# MobiLab – A Mobile Laboratory for On-Site Listening Experiments in Virtual Acoustic Environments

**DOI:** 10.1101/625962

**Authors:** Florian Pausch, Janina Fels

## Abstract

Research in the last decades has pointed out negative effects of noise on cognitive processing and general health. Depending on the frequency and level of exposure as well as individual shielding potential, risk groups like children or elderly people may be particularly vulnerable to commonly known consequences including a reduced attention span, influencing the learning success during education, increased blood pressure and cardiovascular problems or sleep disturbances. To better understand the susceptibility to noise, specifically designed listening experiments in controlled laboratory environments are indispensable but might put high demands on older study participants whose travel expenses to test locations could increase due to reduced mobility. For children, organizational issues like insured transport and supervision during waiting periods, necessitating additional resources, need to be resolved. In order to facilitate practical and efficient study execution, we therefore implemented a mobile hearing laboratory by acoustically optimizing the interior of a caravan. All necessary technical facilities were integrated to perform listening experiments in virtual acoustic environments under controlled conditions directly on site, for example, in front of schools or senior residential centers. The design and construction of this laboratory are presented here and evaluated through acoustic measurements.

## 1 Introduction

In the scope of sustainable office building indoor environment design, auditory distraction appeared as a negative feature in content analysis of sensitive workplaces like open plan offices [1]. In this context, the *irrelevant sound effect*, describing the potential of irrelevant sound to corrupt or disrupt recall of visually presented information, as required in serial recall tasks [2], plays an important role, whereby higher working memory capacity does not necessarily improve shielding [3]. Results of studies investigating the impact of noise on cognitive performance further suggest increased susceptibility of children compared to adults, whether it concerns effects related to indoor noise and reverberation or harmful exposure to aircraft noise [4]. Noise pollution subsequently spans a lifetime and often leads to disabling noise-induced hearing loss among elderly people [5], which might be aggravated by recruitment, distortion, hyperacusis or tinnitus [6], [7]. When conducting specifically designed experiments to investigate noise effects on the mentioned risk groups, controlling extraneous and confounding variables to ensure internal validity of results is one major aspect and becomes particularly challenging when conducting field studies [8]. Here, the application of virtual acoustic environments (VAEs) with full control over virtual sound sources, playing back definable natural source signals, such as speech, music or machine noise, or synthesized noise signals, can be considered as a flexible tool [9]. The validity of this approach is further increased by integration of according source directivities [10], [11], generic or individual receiver directivities [12]–[14], simulation of room acoustics [15] and Doppler shifts in case of moving sources or receivers [16]. Playback of such VAEs can be realized by utilizing techniques that rely on binaural technology, using a pair of headphones for a set of loudspeakers [17], or, if space permits, more hardware-intense loudspeaker-based sound field synthesis methods such as Ambisonics [18] and Wave Field Synthesis [19], [20].

The conduction of listening experiments is usually associated with several additional tasks, like obtaining an ethical proposal or recruiting participants who ideally represent a homogeneous sample, making the procedure a very time consuming one. Increased organizational effort applies in particular to experiments that involve young participants who require the consent of teachers, parents or caregivers and need insured transport and supervision, or elderly participants who might have mobility issues, making the journey to test locations complicated. The idea of bringing the laboratory closer to study participants is imminent and has been realized before for audiometric investigations [21]. In a study by Lankford, Perrone, and Thunder (1999) [22], several mobile audiometric assessment units were checked for compliance with relevant standards [23], [24], pointing out possible inaccuracies in audiometric measurements at certain test frequencies because of exceeded maximum permissible ambient noise levels (MPANLs). The provision of sufficient isolation, particularly in the lower frequency range, can actually be considered one of the biggest challenges in constructing such a mobile test unit.

This article presents another implementation of a mobile laboratory in form of a rebuilt trailer to be used for audiometric testing and the conduction of on-site listening experiments. First, general requirements regarding acoustic and environmental conditions are described. Based on these requirements, design aspects and their implementation are presented in detail. The evaluation of the construction is based on the sound reduction index (SRI) and selected room acoustic (RA) parameters. The results of these measurements will be discussed and compared to normative demands, if applicable, to finally draw conclusions.

## 2 General Requirements

Prerequisites for listening experiments are sufficiently optimized acoustic conditions with decoupling from adjoining spaces to shield low-frequency vibrations, suitable insulation to minimize ambient noise levels (ANLs), low reverberation times, and high speech intelligibility. Hearing booths permanently installed at research institutions, hospitals or worksites, typically fulfill these requirements by meeting the normative requirements for audiometry test rooms [25]– [27]. Transferring these acoustic requirements to a mobile laboratory housed in a caravan with restricted space may, however, complicates the design considerably and might entail compromises, especially in the low-frequency range.

Apart from the acoustic optimization, it is necessary to make the laboratory’s environmental conditions as pleasant as possible. This involves proper seating for the participant in a test room with minimized visual distraction potential, necessitating discrete installation of technical facilities and modest colors of surface materials. Maintenance of a comfortable room temperature, according to local regulations, with the help of a heater or air conditioner (AC) and providing enough fresh air are equally important factors. Only if the ambient conditions are stable across participants can we neglect their influence on experimental results owing to malaise. When controlling the experiment from outside the test room, visual supervision through a window or camera and verbal communication via talkback system must be available, all integrated in an ergonomic workplace [25]. Moreover, security aspects should not be ignored, including the installation of a fire extinguisher and first-aid kit, as well as fluorescent labelling of light switches and standardized emergency exit signs.

## 3 Design and Implementation

### 3.1 General Design

The use of a trailer (Deseo Transport Plus, Knaus Tabbert GmbH, Jandelsbrunn, Germany), see Figure 1 and 3, with increase of maximum load to 1,800 kg, was preferred to that of a motorized mobile home above all for the costs of acquisition, maintenance and non-permanent use. After removing not needed elements, the interior space was divided into test and control room, as depicted in Figure 1b. This division was partly determined by the location of stable attachment points on the caravan’s outer shell, which were required for the acoustic optimization measures described in the following section.

**Figure 1:**
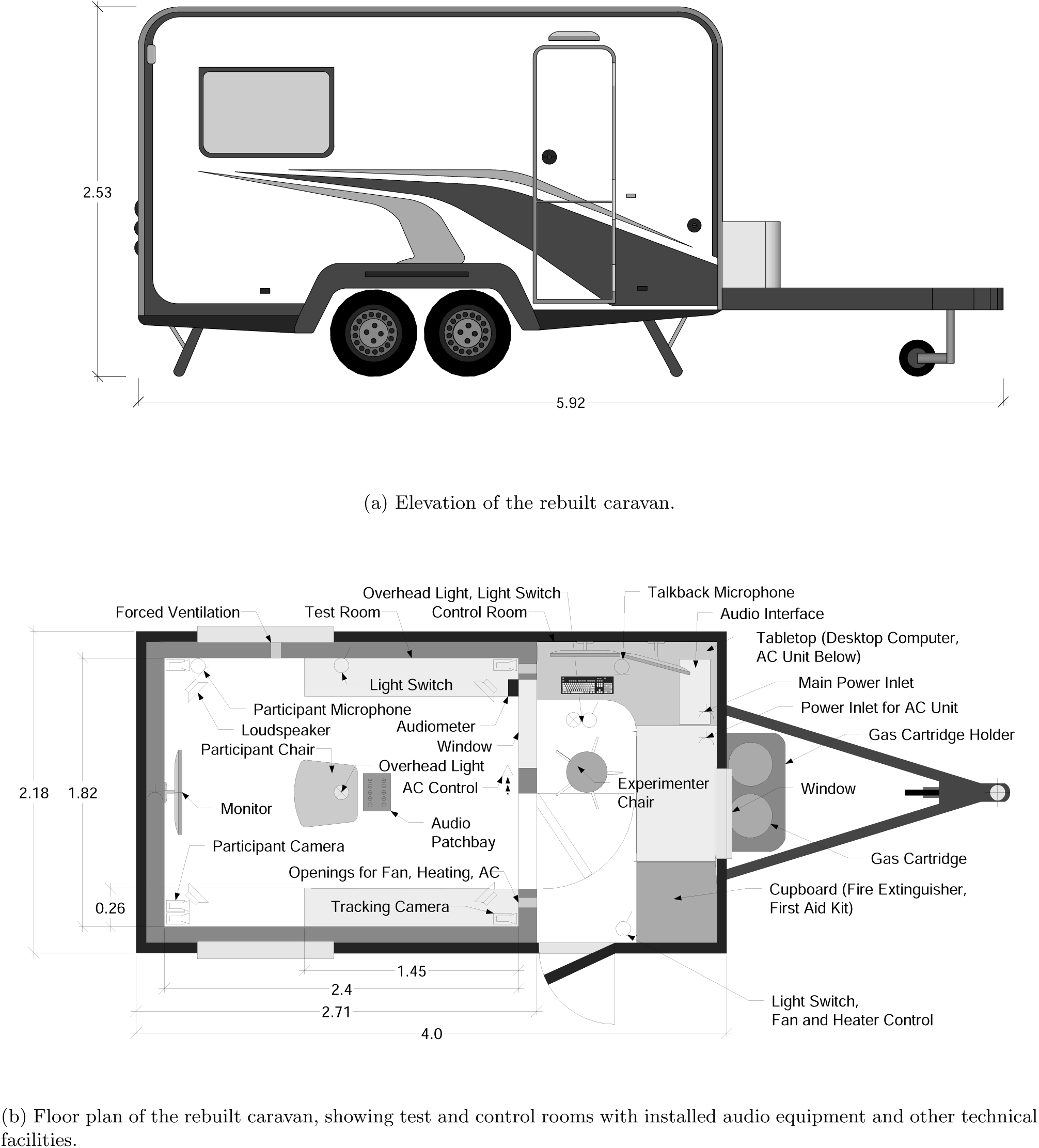
Standard views of the rebuilt caravan used as a mobile laboratory to conduct on-site listening experiments in virtual acoustic environments. Dimensions are given in meters.

### 3.2 Acoustic Optimization

The acoustic optimization strategy can be subdivided into three main measures: (1) Elastic isolation using a low-tuned spring-mass system, commonly referred to as room-within-room construction, (2) adding insulation material between the outer shell and inner room, and (3) placing broadband absorber panels on the test room’s walls and ceiling to minimize reflections in a wide frequency range. Constructional details are shown as cross sections of walls and floor in Figure 2.

**Figure 2:**
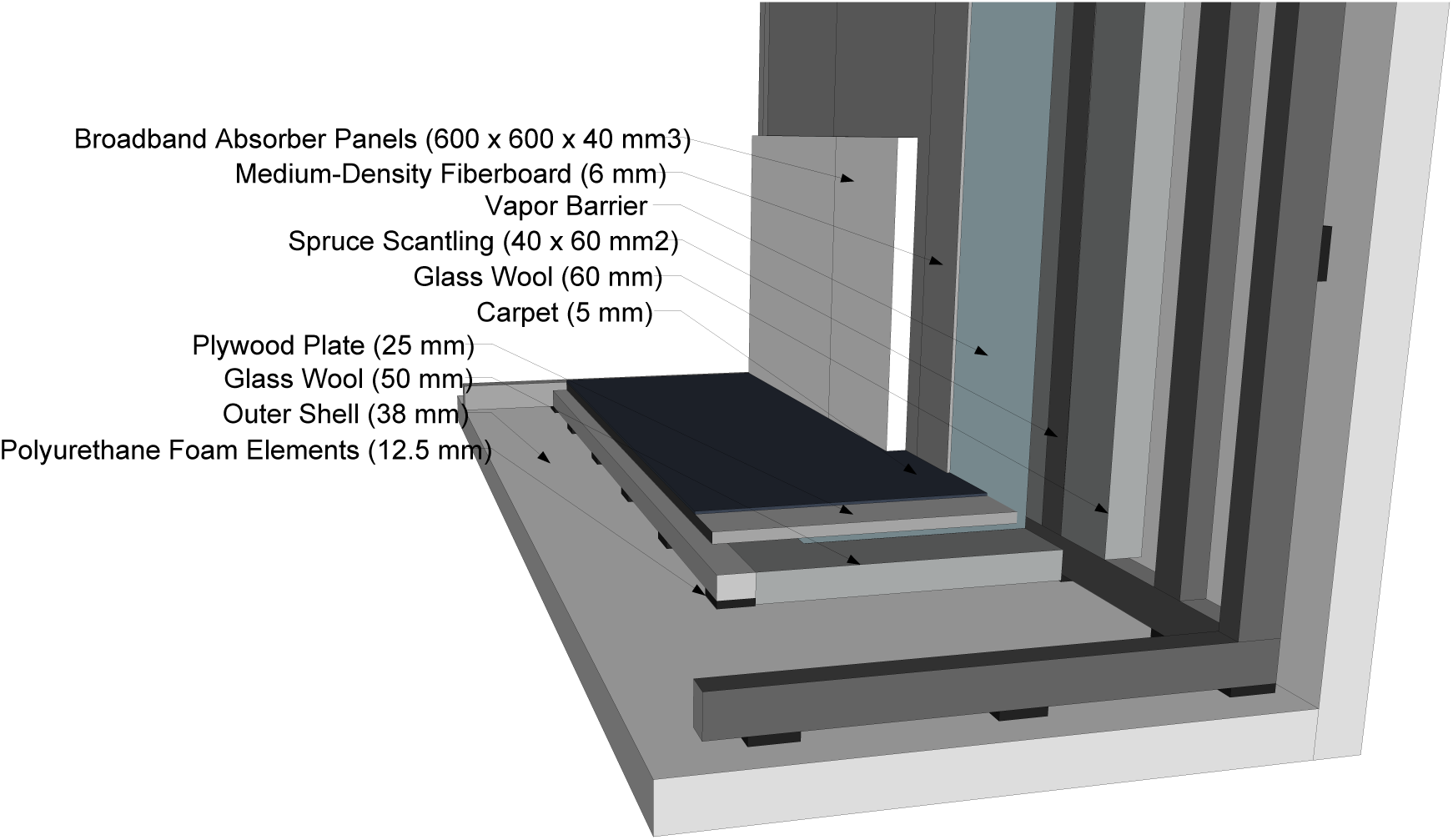
Cross section of floor and walls of the rebuilt caravan’s test room with constructional details.

#### 3.2.1 Elastic Isolation

As supporting element of the test room, a wooden frame made of 40 × 60 mm^2^ (W × D) spruce beams, interlinked with angle irons, was constructed. A door of 610 × 1770 × 39 mm^3^ (W × H × D) was integrated into the frame construction. The frame itself was placed on polyurethane foam elements (Sylomer^®^, Getzner Werkstoffe GmbH, Bürs, Austria) which were selected according to the following considerations, taking into account contact area and static load pressure [28]. Note that the estimations assume a rigid foundation which can only be provided partially by using the caravan supports, cf. Figure1a. Including the wooden frame construction plus further optimization measures (488.5 kg), door and window (36.5 kg), technical equipment (25 kg) and an adult participant (70 kg), we estimated a maximum total weight of approximately 620 kg with a load distribution as outlined in Table1, requiring different types of Sylomer ^®^ elements (12.5 mm thickness).

**Table 1:** Selection of suitable Sylomer^®^ elements based on distributed load and contact areas.

| Area description | Load Distribution (%) | (kg) | Surface Pressure (N/mm<sup>2</sup>) | Static Sinking (mm) | Suitable Sylomer Element |
| --- | --- | --- | --- | --- | --- |
| Walls w/o door and window | 63 | 387.7 | 0.035 | 1.6 | SR42 |
| Wall with door and window | 21 | 129.2 | 0.035 | 1.6 | SR42 |
| Remaining floor area | 17 | 103.4 | 0.023 | 1.7 | SR28 |

A notable mass increase was achieved by inserting a heavy bottom plate (25 mm thick plywood), lowering the resonance frequency *ω*_0_ of the springmass system. Calculating the respective spring stiffnesses of the polyurethane foam elements, depending on pre-defined contact areas, and according partial masses, we estimated the resonance frequencies of the walls to be approximately 18 Hz (26 Hz effectively) and 19 Hz (27 Hz effectively), respectively. For *ω*_0_ *« ω* and *ω « ω*_0_*/η*, with loss factor *η*, the insertion insulation level, given a stiff foundation, would theoretically increase by 12 dB/octave, before decreasing to 6 dB/octave for *ω » ω*_0_*/η* and *ω » ω*_0_ [28]., however, not be achieved given a slightly flexible foundation. In the final implementation, the selected Sylomer^®^ SR42 and SR28 elements covered areas of 0.144 m^2^ and 0.043 m^2^, respectively, corresponding to 19.2 % and 5.2 % of the total floor area of the wooden frame construction, which entails static sinking values as given in Table1. For decoupling at the vertical attachment points and increased safety under dynamic stress during transfer, the wooden frame was additionally connected with the outer shell by custom-made low-vibration bolted joints constructions using Sylomer^®^ SR11 elements.

#### 3.2.2 Insulation

The outer shell of the caravan is a self-sustained full-sandwich construction with maxi-polystyrene isolation, exhibiting wall and floor thicknesses of 31 mm and 38 mm, respectively. Spaces between the wooden frame and outer shell were filled with 60 mm glass wool insulation material (Akustic TP 1, Saint-Gobain Isover, Ludwigshafen, Germany), whereas we only used 50 mm insulation layers between floor and ceiling, aiming at a compromise between room height and insulation performance. To avoid mold formation, a vapor barrier with humidity-variable climate membrane (Vario^®^ KM, Saint-Gobain Isover, Ludwigshafen, Germany) was added. Combining the outer shell with the 6 mm medium-density fiber (MDF) plates, filled with the glass wool layer, we expected sound reduction performance between single and double-leaf constructions.

#### 3.2.3 Absorption

Room acoustic conditions with very low reverberation times can be achieved by full-coverage placement of absorbers on ceiling and walls, including the wheel housings. For this purpose, we used 600 × 600 × 400 mm^3^ (L × W × D) glass wool broadband absorber panels (Saint-Gobain Ecophon, Super G™ B) that are non-flammable, impact-resistant, easy to clean and class-A compliant with French regulations on volatile organic compound (VOC) emissions. According to the product sheet, these absorbers achieve practical absorption coefficients *α*_p_ between 0.25 and 0.8 for octave bands with center frequencies between 125 Hz and 250 Hz, before reaching an *α*_p_ of 1 for octave bands with center frequencies above. We additionally covered the floor with a 5 mm thick fleece carpet. The final inner dimensions of the test room after all optimization measures were 2.4 × 1.82 × 1.77 m^3^ (L × W × H), whereby the covered wheel housings reduced the volume by 2 × (1.45 × 0.26 × 0.28) m^3^ (W × H × D), resulting in a total volume of about 7.5 m^3^, cf. Figure 3c and 3d.

**Figure 3:**
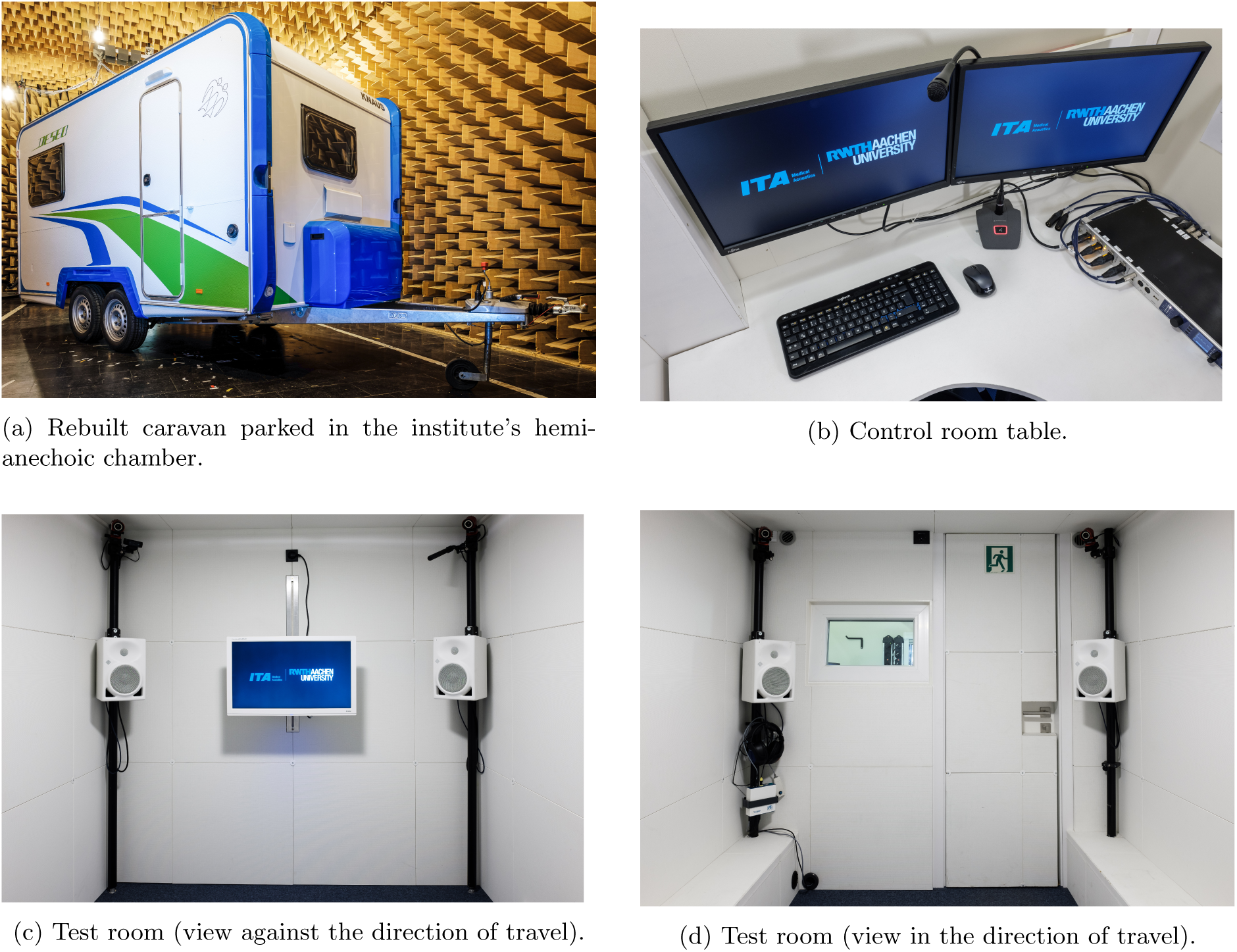
Photos of the implemented mobile laboratory, showing parts of the audio equipment and technical facilities installed in control and test rooms.

### 3.3 Environmental Control

For all-season usage, the caravan was equipped with a sufficiently powered, programmable AC with remote control (Saphir compact, Truma Gerätetechnik GmbH & Co. KG, Putzbrunn, Germany), that additionally dries and cleans the air with the help of fluff and particle filters. For heating purposes, a gas heating (E2400 with fan, Truma Gerätetechnik GmbH & Co. KG, Putzbrunn, Germany) is available. Fresh air supply is provided by the integrated ventilation system. Both test and control room have openings or supply tubes to the mentioned ambient control devices.

### 3.4 Other Technical Facilities

As depicted in Figure 1b, different equipment is necessary to efficiently conduct experiments directly on site. For graphical representation of experimental contents, a 24-inch monitor (Prolite B2480HS, liyama, Tokio, Japan) was attached to a height-adjustable swivel-type mount in the test room. The cushioned participant seat was equipped with a detachable desk which can hold a wireless keyboard (K360, Logitech, Lausanne, Switzerland) and a mouse (M325, Logitech, Lausanne, Switzerland).

#### 3.4.1 Power Supply

The installation of batteries has been deliberately omitted so that an external power connection is required, which is already available at test sites. To avoid exceeding maximum inrush currents when switching on all devices simultaneously via a power strip, it was necessary to provide two separate power supplies, one dedicated solely for the AC.

#### 3.4.2 Audio Devices

Usually, listening experiments include audiometric screening, necessitating a conformal testing device. The installed diagnostic audiometer (Ear 3.0, AURITEC Medizindiagnostische Systeme GmbH, Hamburg, Germany), cf. Figure 3d, is a Class II device for pure tone audiometry, covering a frequency range of 125–16000 Hz, and fulfilling DIN 60645-1:2015-11 (2015) [29]. A maintenance contract ensures regular calibration [25].

Audio signals are controlled via an audio interface (Fireface UFX II, RME Audio AG, Haimhausen, Germany) which is connected to a desktop computer (Intel ^®^ CoreTM i7-7700, 7th generation, 8 GB DDR4 RAM, 2400 MHz, Wi-Fi adapter), both installed in the control room. Two 21.5-inch computer displays (E22-8 TS Pro, Fujitsu, Tokio, Japan) allow for seamless monitoring of the experiment process, see Figure 3b. The sufficiently high number of input channels of the audio interface makes it unnecessary to reconnect microphones that might be needed to measure, for example, the participant’s individual headphone transfer functions [30], [31] or log ANLs in the test room during experiments. Both tasks get additionally simplified by using the audio patchbay, installed directly above the participant, avoiding impractical cable lengths and trip hazards. Playback of simulated signals for the creation of VAEs can be done via a pair of headphones (HD-650, Sennheiser GmbH & Co. KG, Wedemark, Germany) or a set of four height-adjustable loudspeakers (KH 120 AW, Neumann GmbH, Berlin, Germany), see Figure 3c and 3d, that are either fed discretely, by using panning methods for the creation of phantom sources between the physical positions, or by applying more advanced methods like acoustic crosstalk cancellation [17], allowing for three-dimensional virtual sound source placement and room acoustic simulations [15], [32]. Additional output channels for the integration of other playback devices such as research hearing aids [33] are also available through the audio patchbay.

#### 3.4.3 Motion Tracking

Logging and evaluating the participant’s head movements during experiments could provide significant hints to hearing strategies in complex auditory scenes [34]. When playing back VAEs via headphones or loudspeakers with acoustic crosstalk cancellation filters such motion tracking is vital and enables dynamic reproduction that is adjusted in real-time based on the participant’s current head position and orientation, further approaching the simulated real-life scenario [35]. For increased freedom of movement, an optical motion tracking system (OptiTrack, Natural-Point Inc., Corvallis, OR, USA) with four infrared cameras (Flex 13) has been installed, see Figure 3c and 3d, forwarding tracking data to a hub (Opti-Hub 2) that is connected via USB cable to the desktop computer.

#### 3.4.4 Supervision

Visual supervision is made possible by a 600 × 400 mm^2^ (W × H) window (IDEAL 4000, 5-chamber system with two seals, triple glazing, Neuffer Fenster+Türen GmbH, Stuttgart, Germany), installed inclined and pointing towards the ceiling to prevent unfavorable reflections at the participant’s listening position, see Figure 3d. We additionally integrated a web cam (C920 HD Pro Webcam, Logitech, Lausanne, Switzerland) in the left upper room corner with respect to the participant’s viewing direction, see Figure 3c.

Providing oral instructions before and answering questions or evaluating responses by the participant during the experiments can be done via the available intercom system, consisting of a participant microphone (NTG2, Røde Pvt. Ltd., Sydney, Australia) and a control room microphone (DMG-700, Monacor, Bremen, Germany) with table socket (MAT 133-S B, Sennheiser GmbH & Co. KG, Wedemark, Germany). The respective interlocutor signal is either played back via headphones or loudspeakers in the test room, and headphones (HD-650, Sennheiser GmbH & Co. KG, Wedemark, Germany) in the control room, cf. Figure 3b–3d.

#### 3.4.5 Security Aspects

In case of unforeseen events, like power failure or panic attacks by the participant, both test room exit and main exit are indicated by fluorescent signs [36] with additional marking of the doorstep by a warning stripe. The light switch in the test room also carries a fluorescent sticker and is in tangible proximity for the participant. For the treatment of injuries and in the event of fire, the cupboard in the control room contains a first aid kit [37] and a 1-kg fire extinguisher [38], respectively, both indicated by according normative stickers [36]. Safety regulations for the use of gas dictate forced ventilation by means of an air outlet installed in one of the test room’s bottom edges, cf. Figure 1b.

## 4 Methods

### 4.1 Sound Reduction Index

Audiometric standards demand MPANLs which are depending on the type of audiometric screening [25]–[27]. For on-site conduction of listening experiments, it is of particular interest how much exterior noise may be present without distorting headphone-based audiometric screening results and the experiment. To what extent these requirements are met by the implemented mobile laboratory shall be checked in the light of measurement results.

The SRI can be considered as a suitable metric to quantify the frequency-dependent insulation performance of the implemented mobile laboratory. Instead of diffuse sound field conditions, ISO 3744:2010 [39] allows sound source power determination under free-field conditions using measurement points distributed on an enveloping hull above a reflecting surface. We therefore parked the caravan in the institute’s hemi-anechoic chamber with dimensions 11 × 5.97 × 4.5 m^3^ (L × W × H), exhibiting a lower frequency limit of about 100 Hz, and determined the SRI indirectly via sound power level (SWL) measurements in two consecutive measurement cycles. As omnidirectional sound source we used a three-way dodecahedron loudspeaker, developed at the Institute of Technical Acoustics, RWTH Aachen University [40]. In both cycles, the loudspeaker was placed at a height of 1.45 m, measured from the floor of the hemi-anechoic chamber to the center of the mid-range driver, one time without the caravan and the other time in the center of the caravan’s test room. Exponentially swept sines were generated in MATLAB (The MathWorks, Inc., Natick, MA, USA) at a sampling frequency of 44.1 kHz and a length of 2^20^ samples. The measurement signal was D/A-converted (RME Madiface XT, Audio AG, Haimhausen, Germany), bandpass-filtered as per loudspeaker crossover specifications, energetically band-matched using a digital loudspeaker controller (HD2, FourAudio, Herzogenrath, Germany) and sent through a custom-made Class B power amplifier to be played back by the measurement loudspeaker. Sweep responses were captured by a set of microphones (KE4, Sennheiser GmbH & Co. KG, Wedemark, Germany) that sampled a cuboid hull [39, Figure C.10] at a distance of 1 m to the reference cuboid, set by the outer dimensions of the caravan without drawbar. The calibrated microphone signals were A/D-converted and amplified (RME Octamic XTC, Audio AG, Haimhausen, Germany) before being deconvolved in MATLAB using the ITA-Toolbox [41]. For sufficient signal-to-noise ratio, the measurements were repeated 4 and 16 times in the first and second measurement cycle, respectively. We then calculated the sound pressure levels (SPLs) in third-octave bands at all microphone positions, summed up the weighted SPLs per measurement cuboid sub-area referenced to the measurement cuboid total area, and finally obtained the SWL [39]. The extraneous noise correction factor *K*_1_ has not been applied since signal-to-noise ratios exceeded the minimum value of 15 dB in all third-octave bands. Also, the environmental correction factor *K*_2_ has been omitted as absolute SWLs were not of interest [39]. The SRI was subsequently derived by taking the difference between third-octave band SWLs, obtained in the first and second measurement cycle.

The calculated SRI need to be related to normative requirements of [25] to estimate MPANLs in the test room when parking the caravan at test locations. Previously published traffic noise levels [42] might, however, not be representative as the rebuilt caravan will not be parked close to heavily travelled streets. Instead, we measured unweighted equivalent SPLs LZeq at three typical testing sites for planned future experiments, namely local school yards, for a measurement duration of 15 min each, using a Class I sound level meter (NTi XL2, M2230 microphone, Schaan, Liechtenstein) [43], set at low level measurement range to cover 0 to 100 dB re 20 *µ*Pa. The measurements were carried out during periods in which either children were present on the respective school yard or during teaching hours when no activities took place.

### 4.2 Room Acoustics

As loudspeaker-based playback methods are sensitive to unwanted reflections [19], [44], optimized room acoustics, including low reverberation times and high speech clarity, are crucial for accurately reproducing VAEs and unbiased listening experiments. We therefore conducted room acoustic measurements based on ISO 3382-2 (2008) [45] to quantify the mobile laboratory’s test room in terms of selected standardized parameters. Although special care was taken to optimize the two source and three receiver positions (standard measurement precision), normative requirements concerning minimal distance from walls could not be met given the space available. Additionally, owing to full-coverage placement of absorbers on ceiling and walls, the sound field in the room cannot be declared as diffuse, thus measurement results should be interpreted carefully.

As excitation signal, an exponential sweep with a length of 2^18^ samples at a sampling frequency of 44.1 kHz, covering a frequency range of 20–20000 Hz, was generated in MATLAB, converted from digital to analog domain (RME Fireface UC, Audio AG, Haimhausen, Germany) and further processed and played back as done for the SRI measurements. For recording the sweep responses, we used a ½” diffusefield compensated microphone (Type 4134, Brüel & Kjær, Nærum, Denmark) with microphone preamplifier (Type 2669, Brüel & Kjær, Nærum, Denmark) and microphone conditioner (Type 2690, Brüel & Kjær, Nærum, Denmark). Room acoustic parameters *T*_30_ and *C*_50_ were calculated per source-receiver combination with MATLAB and the ITA-Toolbox [41], applying noise detection and compensation [46], with subsequent energetic averaging over spatial measurement results per octave band.

## 5 Results

### 5.1 Sound Reduction Index

Measured unweighted SWLs in third-octave bands of the dodecahedron loudspeaker, obtained in the two measurement cycles, are plotted in Figure 4. Under free-field conditions, the overall SWL exhibits a value of 92 dB(Z) for the playback level set, with values within ±4 dB around a mean value of 78 dB(Z) in the frequency bands with center frequencies between 50 Hz and 16 kHz. In the second measurement cycle, SWL values showed a high-frequency sloping, owing to the insulation properties of the test room construction with an overall SWL of 69 dB(Z).

**Figure 4:**
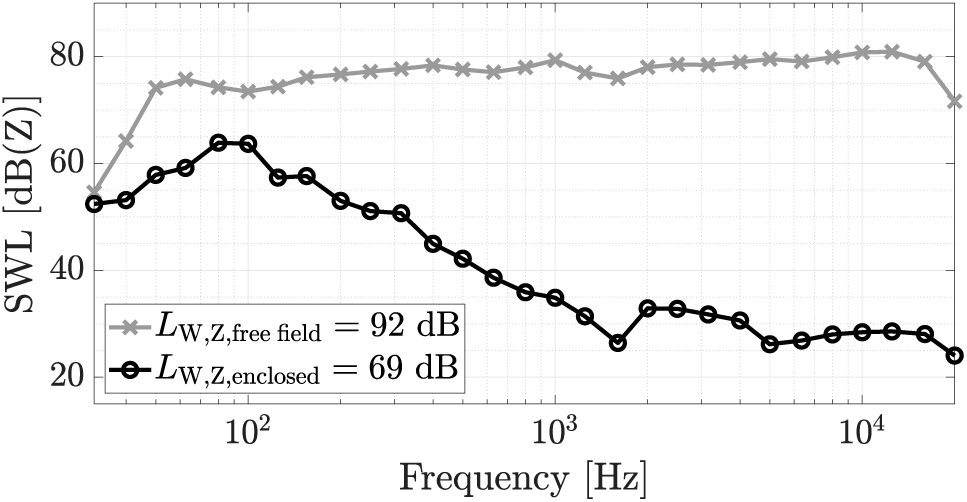
Unweighted sound power level (SWL) of the dodecahedron loudspeaker in third-octave bands, measured under free-field conditions above a reflecting surface (solid gray line with crosses), *L*_W,Z,free field_, and in the caravan’s test room (solid black line with circles), *L*_W,Z,enclosed_, resulting in overall SWLs of 92 dB(Z) and 69 dB(Z), respectively.

These insulation properties can be better analyzed by looking at the SRI, plotted in Figure 5. The insulation performance lies between single and double leaf construction and shows an increase of 9 dB/octave in SRI above the resonance frequency around 90 Hz. To obtain a representative single integer number, the measurement results are compared with the shifted reference curve defined in ISO 717-1:2013 (2013) [47], resulting in a weighted difference level of 35 dB at 500 Hz.

**Figure 5:**
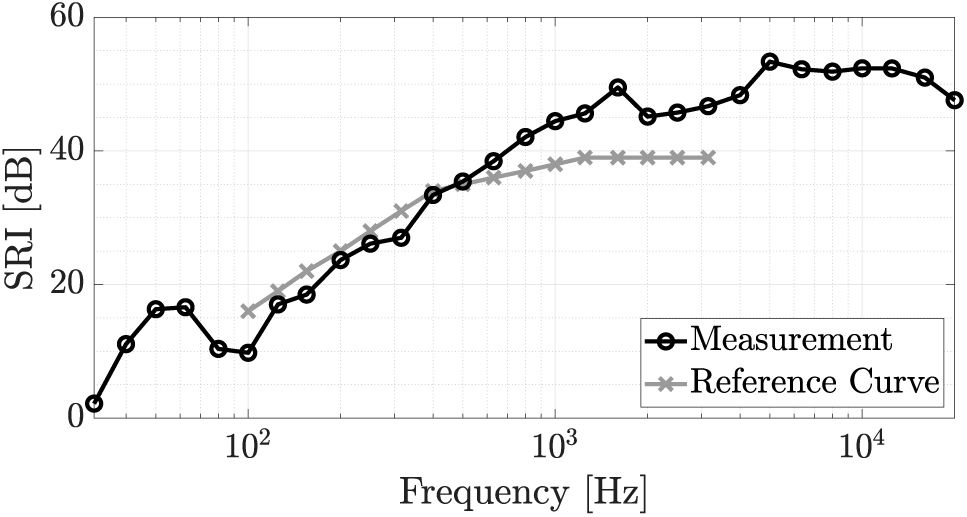
Sound reduction index (SRI, solid black line with circles) calculated as difference between sound power levels LW, Z, free field and LW, Z, enclosed in third-octave bands. Shifting the reference curve [47] (solid grey line with crosses) resulted in a weighted difference level of 35 dB at 500 Hz.

The corrected measurement results of unweighted equivalent SPLs LZeq after subtracting the obtained SRI values per third-octave band, measured at the three representative future test locations, are shown in Figure 6. Plotted alongside the MPANLs as per DIN ISO 8253-1:2011-04 (2011) [25] for air conduction audiometry (gray bars), using a pair of normative supra aural headphones for measuring hearing levels down to 0 dB, the results provide an estimate of practicability. In case of activity on the school yard, these thresholds are exceeded for third-octave bands with center frequencies below 1 kHz (dotted gray line) while being maintained for frequency bands above (solid gray line). No school yard activity, on the other hand, resulted in an estimated interior noise floor complying to MPANLs in a wide frequency range (solid black line), while only exceeding the limits in third-octave bands with center frequencies between 80 and 200 Hz (dotted black line).

**Figure 6:**
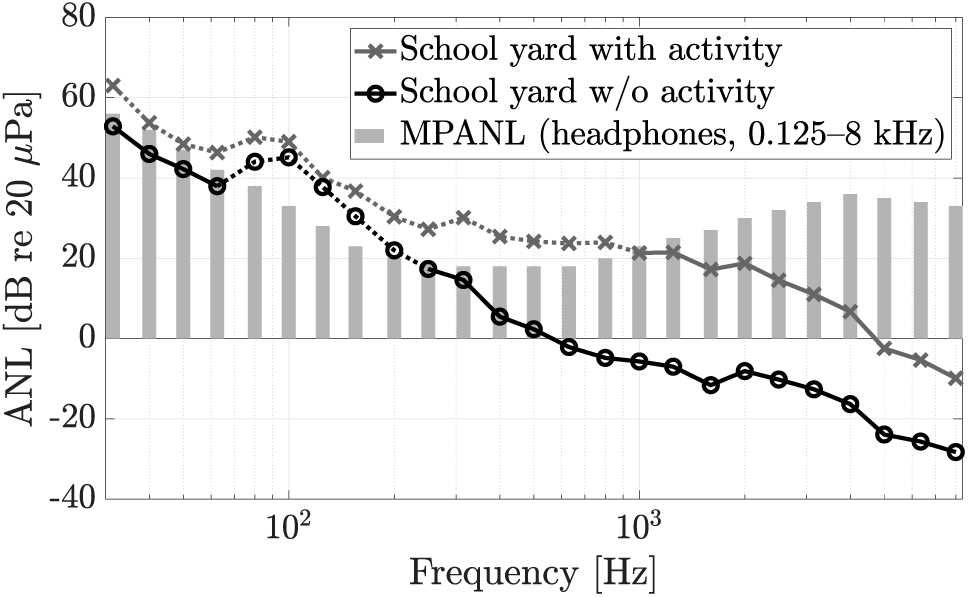
Ambient noise levels (ANLs) to be expected in the mobile laboratory’s test room. Solid and dotted gray and black lines with crosses and circles represent mean equivalent sound pressure levels (SPLs) LZeq in third-octave bands measured in-situ at three representative school yards with and without activity, respectively, after subtracting sound reduction indices (SRIs). Results are compared to maximum permissible ambient noise levels (MPANLs) as per DIN ISO 8253-1:2011-04 (2011) [25] when using supra-aural headphones and audiometric test frequencies between 125 and 8000 Hz (grey bars).

### 5.2 Room Acoustics

Results of mean reverberation times *T*_30_ in octave bands are plotted in Figure 7, showing an increase towards lower and higher frequencies while in general exhibiting very low values in the considered frequency range with a mean mid-frequency value of 50 ms (averaged between the values of the octave bands with center frequencies of 500 Hz and 1 kHz).

**Figure 7:**
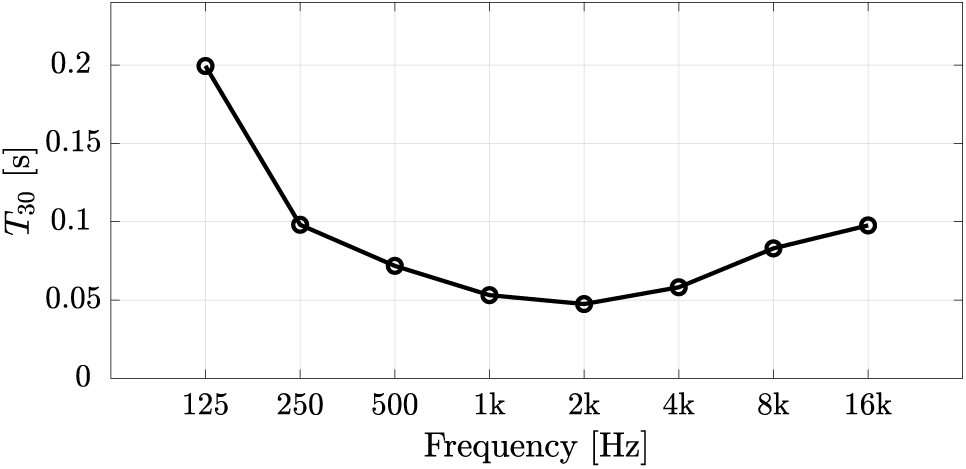
Mean reverberation times *T*_30_ of the mobile laboratory’s test room in octave bands.

Speech clarity values *C*_50_, as plotted in Figure 8, are particularly high and result in an intelligibility-weighted and summed single value of 54 dB, predicting excellent speech clarity [48].

**Figure 8:**
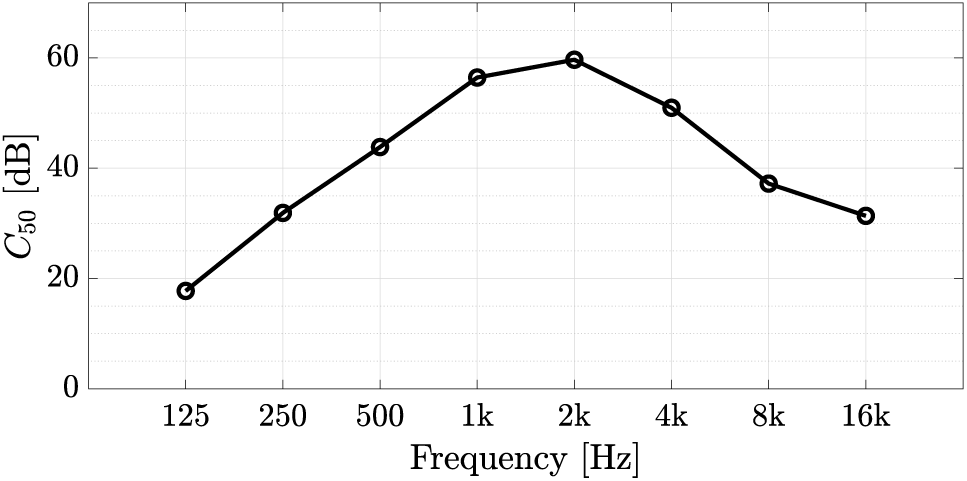
Mean speech clarity values *C*_50_ of the mobile laboratory’s test room in octave bands.

## 6 Discussion

### 6.1 Sound Reduction Index

Owing to the medium-heavy construction, the sound reduction performance in lower frequencies exhibits merely low values. If the test room was built exclusively for the conduction of audiometries, the integration of a heavily damped room-within-room sound cabin, installed additionally in the test room, would have substantially improved low frequency insulation performance [21]. This construction possibility was knowingly neglected to meet the additional spatial requirements for loudspeaker-based listening experiments or audiometries, the latter demanding a minimum loudspeaker distance of 1 m [26]. Interestingly, the achieved insulation performance is very similar to the one reported by Taylor, Burns, and Mair (1964) [21] for the vehicle shell alone. Given the estimated ANLs in the test room in case of additional activities on the school yard, the implemented mobile laboratory does not provide sufficient isolation performance at lower frequencies to rule out a potential bias in audiometric and experimental results and therefore should only be operated during teaching hours, when there is no activity on the school yard.

Speaking of audiometric results, participants involved in experiments following standard pure tone audiometric screening must have hearing capacities above a predefined threshold across all test frequencies to be admitted, which typically corresponds to 15–20 dB hearing level for normal hearing. Apart from isolated low-frequency sensorineural hearing loss, the insulation properties in presence of reasonably low external noise enable proper detection of other types of sensorineural hearing loss, usually indicated by audiograms with high-frequency sloping, or conductive hearing loss, for test frequencies of 250 Hz and above. For the purpose of documentation, reproducibility and validity of results, it is nevertheless recommended to measure any ambient noise present during audiometry or the experiment by means of an additional measurement microphone installed in the caravan’s test room, which is feasible at least for headphone-based listening experiments. The conduction of more specialized studies, involving participants with hearing loss, usually demands audiometries that were anyway performed in fully standard-compliant hearing booths prior to the actual experiment, allowing for accurate participant classification or hearing aid fitting when using a master hearing aid platform [33], [49], [50].

### 6.2 Room Acoustics

The two evaluated parameters *T*_30_ and *C*_50_ both reflect excellent conditions of the test room for conducting loudspeaker-based listening experiments with minimized detrimental effects of additional reflections [44]. For loudspeaker-based binaural reproduction using acoustic crosstalk cancellation, channel separation is among the most important reproduction quality metrics. As the reverberation times measured in the test room are even lower than those reported by Pausch, Aspöck, Vorländer, *et al.* (2018) [33], the practically achievable channel separation will be at least comparable or potentially higher, typically increasing towards higher frequencies, while exceeding the required minimum for sufficient binaural signal perception [51]. Furthermore, experiments under free-field conditions or with simulated room acoustics are minimally distorted by the listening environment.

## 7 Summary and Conclusion

The design and construction of a mobile laboratory has been presented and evaluated through acoustic measurements. Equipped with all necessary technical equipment, including devices to maintain pleasant ambient conditions, the refurbished caravan can be used for an efficient on-site conduction of audiometries and listening experiments when parked at test locations with ambient noise levels that are reasonably low in relation to the achievable insulation properties of the construction. Subject to optimized room acoustics in terms of very low reverberation times and excellent speech intelligibility, the laboratory’s test room allow the reproduction of simulated acoustic test scenarios to assess, for example, speech and language understanding under challenging room acoustic conditions and the impact of additional noise sources in particularly vulnerable investigation groups, using controlled and non-intrusive virtual acoustic environments.

## Acknowledgements

We want to thank the team of our mechanical workshop, Uwe Schlömer, Thomas Schäfer and Marc Eiker, for the design implementation as well as Rolf Kaldenbach for planning and conducting electrical installations. Additional thanks go to Mark Müller-Giebeler and Gottfried Behler for helpful advice and assistance during the design process and the measurements, Philipp Seltsam for conducting the on-site sound level measurements, Hark Braren for critical comments, and Karin Charlier for handling financial matters.

